# Voxel-to-voxel predictive models reveal unexpected structure in unexplained variance

**DOI:** 10.1101/692319

**Authors:** Maggie Mae Mell, Ghislain St-Yves, Thomas Naselaris

## Abstract

Encoding models based on deep convolutional neural networks (DCNN) more accurately predict BOLD responses to natural scenes in the visual system than any other currently available model. However, DCNN-based encoding models fail to predict a significant amount of variance in the activity of most voxels in all visual areas. This failure could reflect limitations in the data (e.g., a noise ceiling), or could reflect limitations of the DCNN as a model of computation in the brain. Understanding the source and structure of the unexplained variance could therefore provide helpful clues for improving models of brain computation. Here, we characterize the structure of the variance that DCNN-based encoding models cannot explain. Using a publicly available dataset of BOLD responses to natural scenes, we determined if the source of unexplained variance was shared across voxels, individual brains, retinotopic locations, and hierarchically distant visual brain areas. We answered these questions using voxel-to-voxel (vox2vox) models that predict activity in a target voxel given activity in a population of source voxels. We found that simple linear vox2vox models increased within-subject prediction accuracy over DCNN-based models for any pair of source/target visual areas, clearly demonstrating that the source of unexplained variance is widely shared within and across visual brain areas. However, vox2vox models were not more accurate than DCNN-based models when source and target voxels came from separate brains, demonstrating that the source of unexplained variance was not shared across brains. Furthermore, the weights of these vox2vox models permitted explicit readout of the receptive field location of target voxels, demonstrating that the source of unexplained variance induces correlations primarily between the activities of voxels with overlapping receptive fields. Finally, we found that vox2vox model prediction accuracy was heavily dependent upon the signed hierarchical distance between the source and target voxels: for feed-forward models (source area lower in the visual hierarchy than target area) prediction accuracy decreased with hierarchical distance between source and target. It did not decrease for feedback models. In contrast, the same analysis applied across layers of a DCNN did not reveal this feed-forward/feedback asymmetry. Given these results, we argue that the structured variance unexplained by DCNN-based encoding models is unlikely to be entirely caused by spatially correlated noise or eye movements; rather, our results point to a need for brain models that include endogenous dynamics and a pattern of connectivity that is not strictly feed-forward.

## 1. Introduction

A critical measure of understanding in visual neuroscience is the ability to predict how the brain will respond to arbitrary, complex stimuli. Models that predict brain activity in response to visual stimuli are known as encoding models [1, 2, 3, 4, 5]. Currently, the most accurate encoding models for predicting responses to natural scene stimuli in visual cortical areas are based upon deep convolutional neural networks (DCNN) that have been trained on object recognition tasks [5, 6, 7, 8].

DCNN-based encoding models are impressive in their ability to accurately predict BOLD activation measured in voxels across the hierarchy of visual areas using a single underlying model of computation. Nonetheless, DCNN-based encoding models (and, to our knowledge, all encoding models) fail to accurately predict brain activity in response to natural scenes in *most* voxels in all visual areas (Fig. 1A, left). Intriguingly, the voxels for which encoding models make accurate predictions tend to have receptive fields located in the periphery of the visual field, while the voxels for which encoding models fail to make accurate predictions are concentrated about the foveal representation of the retinotopic map in all areas (Fig. 1A, middle and right).

**Figure 1:**
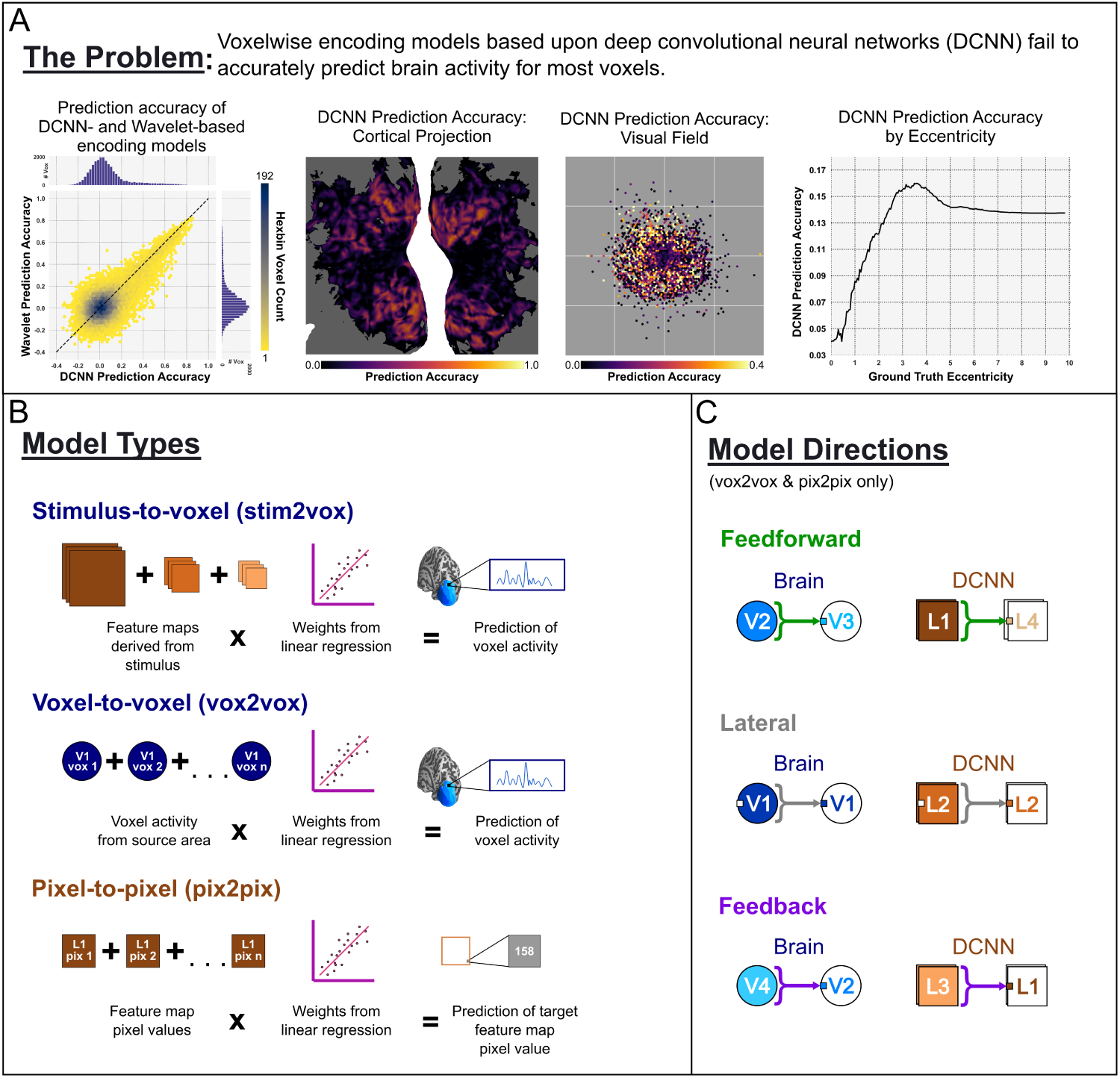
Problem and Approach. **A: The problem** Visualizations of the prediction accuracy of the DCNN-based encoding model. *Left* : The joint distribution of prediction accuracy (Pearson correlation between predicted and measured brain activity) for the DCNN-based encoding model (x-axis) and Gabor wavelet-based encoding model (y-axis; data taken directly from [5]). The slightly higher count (color, yellow=low count, dark blue = high count, white = no data) of voxels below the line at unity (dashed) reveals the advantage of the DCNN-over the wavelet-based encoding model. We refer to the high-count cluster of voxels at the origin as the “ball of nothingness”, since neither model explains any variance for these voxels. *Left Middle*: Prediction accuracy of the DCNN-based encoding model (color) projected onto a cortical flatmap. Prediction accuracy is poorest (dark purple) in the foveal representation. *Right Middle*: Prediction accuracy of the DCNN-based encoding model (color indicates median) projected into visual space (gray square) using the receptive locations (hexagonal bins) of all voxels. Prediction accuracy is poorest for voxels with foveal receptive fields (bins near center of square). *Right* : Prediction accuracy of the DCNN-based encoding model (y-axis indicates median) against receptive field eccentricity (x-axis). **B: Model Types**. *stim2vox* : The DCNN-based encoding model is a stimulus-to-voxel (stim2vox) model that transforms stimuli into a set of feature maps (brown squares) and then into a prediction of voxel activity (blue curve). In the DCNN-based encoding model the transformation of stimuli into feature maps is performed by a deep neural network; the transformation from feature maps to voxel activity is estimated via linear regression (idealized pink line). *vox2vox* : In a voxel-to-voxel (vox2vox) model activity in a population of source voxels (blue circles) is linearly transformed into a prediction of activity in a target voxel. *pix2pix* : In a pixel-to-pixel (pix2pix) model activity in a population of source pixels in a feature map of the DCNN (brown squares) is linearly transformed into a prediction of activity of another target pixel in the DCNN. **C: Model Directions**. *Feed-forward* (shown here and throughout all figures in green) indicates a model that uses data from a lower (i.e., closer to eye or stimulus) source area/layer to predict a target in a higher area/layer. In these examples, brain activity from source voxels in V2 is used to predict activity of one target voxel in V3 (V2 → V3). Pixel values from feature maps in Layer 1 of the DCNN are used to predict the value of one pixel from one feature map in Layer 4 (L1 → L4). *Lateral* (shown in grey) models use data from a source area/layer to predict a target within the same area/layer. For example, V1→V1 or L2→L2. *Feedback* (purple) models use data from a source area/layer to predict a target in a lower area/layer. For example, V4→V2 or L3→L1.

In this paper we investigate the structure of the variance that the DCNN-based encoding model can’t explain. We frame this investigation as a series of questions about the source or sources of the unexplained variance. We first address the question of scale: is there a unique source of unexplained variance that is unique to each voxel, or a common source that is shared across voxels? To answer this question, we applied a voxel-to-voxel (vox2vox) modeling approach (Fig 1B; [9, 10, 11]). Unlike stimulus-to-voxel (stim2vox) encoding models (e.g., the DCNN-based encoding model), vox2vox models use activity in a population of source voxels to predict activity in a target voxel. By fitting vox2vox models for different source/target pairings we can determine the extent to which the variance unexplained by stim2vox models can be explained by the activities of other voxels. If much of the variance unexplained by the stim2vox model can be explained by the vox2vox model, we can infer that the causes of unexplained variance affects both target and source.

We then use the vox2vox encoding models to address more detailed questions about the variance unexplained by stim2vox models. Are the sources of unexplained variance specific to individual subjects, or shared between them? To answer this question we use source voxels in one individual’s brain to predict activity in target voxels from another individual’s brain. Are the sources of unexplained variance independent of or in register with retinotopic representations in cortex? To answer this question we use the vox2vox model weights to read out the receptive field location of target voxels given the receptive field locations of source voxels. Finally, are the sources of unexplained variance shared across hierarchical levels? To answer this question we investigate how the amount of variance that can be explained using vox2vox models depends on the hierarchical relationship between areas. The answers to these questions provide a profile of a potent brain signal that hides in plain sight, is not entirely stimulus-driven but is almost certainly not noise.

## 2. Methods

### 2.1. Data

We analyzed data from two fMRI experiments: a standard retinotopic mapping experiment, and the publicly available vim-1 natural scenes dataset [12]. The vim-1 dataset includes BOLD responses to 1,870 natural scene photographs for two subjects. Voxels were localized to regions of interest (ROI) including V1, V2, V3, V4, V3a, V3b, and LO (for our analyses we combined V3a and V3b into one area, V3ab). The retinotopic mapping experiment featured standard rotating wedge, expanding ring, and drifting bar stimuli. This experiment was completed by subject S1 from the vim-1 dataset. Coverage included all areas named above. All subjects provided informed consent prior to scanning.

### 2.2. Encoding Models

#### 2.2.1. Stimulus-to-Voxel

Two stim2vox models were used to generate ground truth receptive field information and predict voxel activity. For all voxels in the vim-1 natural scenes dataset, the feature-weighted receptive field model (fwRF) was applied to the feature maps of a deep convolutional neural network. Full details regarding this model can be found in St-Yves and Naselaris [5]. Briefly, the fwRF is a form of voxel-wise encoding model that separates the specification and estimation of receptive field location and size from feature tuning. The fwRF uses the following model to generate predictions of brain activity, 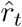, in response to a visual stimulus *S*_*t*_:

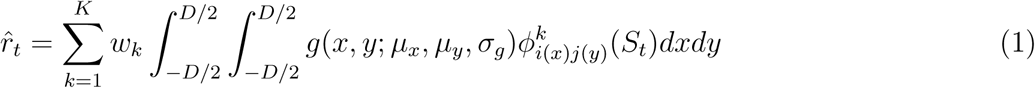

where *D* is the visual angle sustained by the image, the function 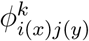 specifies the value of pixel (*i, j*) of the *k*^*th*^ feature map applied to the stimulus *S*_*t*_, and *g*(*x, y*; *μ*_*x*_, *μ*_*y*_, *σ*_*g*_) is the feature pooling field, which is an isotropic-2D Gaussian function, with center (*μ*_*x*_, *μ*_*y*_) and radius *σ*_*g*_. The feature pooling field indicates the region of visual space in which stimulus variation induces variations in activity of the voxel. The feature weights, *w*_*k*_, indicate the features encoded in the activity of the voxel. The set of feature maps used are the same for each voxel, but the weights assigned to each feature will vary. In this paper, the feature weights for the stim2vox model were the feature maps of a DCNN with one input layer, five convolutional layers and three fully-connected layers. This DCNN was trained to classify images in the ImageNet database [13].

The location and radius of the feature-pooling field, as well as the feature weights are estimated by minimizing the sum-of-squared prediction error between model output and brain activity for each voxel over the set of image/response pairs in the training set. Values for the location and radius of the feature-pooling field, i.e. the fwRF center and size, are inferred via a brute force search through a grid of candidate locations and radii. Values for the feature weights are estimated using stochastic gradient descent.

In Figure 1A we compare prediction accuracy of a Gabor wavelet-based model to the prediction of the DCNN-based model. The wavelet-based model was estimated using the same fwRF framework as described above, except that feature maps were constructed by filtering images with a bank of Gabor wavelets. See [5] for complete details.

For retinotopic mapping experiments a population receptive field (pRF) analysis was used. Details of this method can be found in [3] and [14].

#### 2.2.2. Voxel-to-voxel

Voxel-to-voxel models linearly combine activity from one brain area to predict activity in one voxel:

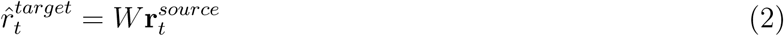

where 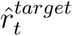 is the predicted activation of the target voxel, *W* is a matrix of vox2vox model weights, and 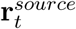 is an array of activations from source voxels. We used ridge regression to determine the weights assigned to each voxel in a source area. We fit separate vox2vox models for each pair of visual areas named above. Thus, for each target voxel we fit six distinct vox2vox models corresponding to the seven ROIs named above.

For each pair of ROI’s we refer to a vox2vox model as “feedforward” if the source voxels are lower in the hierarchy of ROIs than the target voxel. We refer to a vox2vox model as “feedback” if the source voxels are higher in the hierarchy than the target voxel. We refer to a vox2vox model as “lateral” if the source and target voxels are in the same ROI (Fig 1C). The hierarchy of ROIs is defined by the sequence V1, V2, V3, V4, LO/V3ab, where V1 is the “lowest” ROI in the hierarchy.

#### 2.2.3. Pixel-to-pixel

Pixel-to-pixel models (Figure 1B) linearly combine activity in the pixels of one DCNN layer to predict activity of a single target pixel:

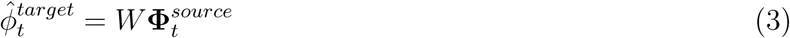

Where 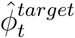 is the activation of a target pixel in one of the feature maps of the DCNN, *W* are the pix2pix model weights, and 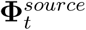 is an array of activations of source pixels taken from all feature maps in one layer of the DCNN. This DCNN was trained to classify images in the CIFAR-10 dataset [15].

Note that pix2pix models ignore the connection weights learned when the DCNN was trained to recognize objects in natural scenes. As with vox2vox models we fit a pix2pix model for every possible pair of source layer and target layer, and refer to pix2pix models as feedforward, feedback and lateral depending upon the relative positions of the source and target layers in the network hierarchy.

### 2.3. Prediction accuracy and cross-validation

All encoding models were trained on 1750 responses to natural scene photographs and tested on the remaining 120. Ridge regression hyper-parameter values were selected via line search by cross-validating against 20% of the training data. Prediction accuracy is the Pearson correlation between model predictions and measured activity (in the brain for vox2vox models, in the DCNN for pix2pix models).

### 2.4. Surface Reconstructions

Cortical reconstruction, segmentation, pial and white-matter surface rendering and cortical inflation of T1-weighted volumes were passed to Freesurfer’s recon-all (v6) [16]. Relaxation cuts on the inflated surface were made using Blender (v2.78) and then imported into pycortex [17] for flattening and final rendering. Functional data were aligned to the structural T1 with FSL FLIRT [18] and projected onto cortical surfaces in pycortex.

## 3. Results

### 3.1. The source of variance unexplained by stim2vox models is shared within and between visual areas

To determine if unexplained variance is driven by a shared source we compared the prediction accuracy of vox2vox models to a DCNN-based encoding model ([5]; hereafter referred to as the “stim2vox” model) for each source-target pairing. Thus, for each source-target pairing, we visualized the joint distribution of stim2vox (y-axis) and vox2vox (x-axis) model prediction accuracy. Even though vox2vox models are linear, vox2vox models have higher cross-validated prediction accuracy than the stim2vox model for nearly every target voxel in every source/target pairing (Fig. 2 left). The most dramatic improvement in prediction accuracy was for the large mass of foveal voxels–described in Fig. 1A as “the ball of nothingness”–for which both the DCNN-based encoding model and its wavelet-based predecessor ([4]) completely failed. Relative to the stim2vox model this improvement homogenizes prediction accuracy across the cortical flatmaps (Fig. 2 middle left) and the visual field (Fig. 2 middle right and right).

**Figure 2:**
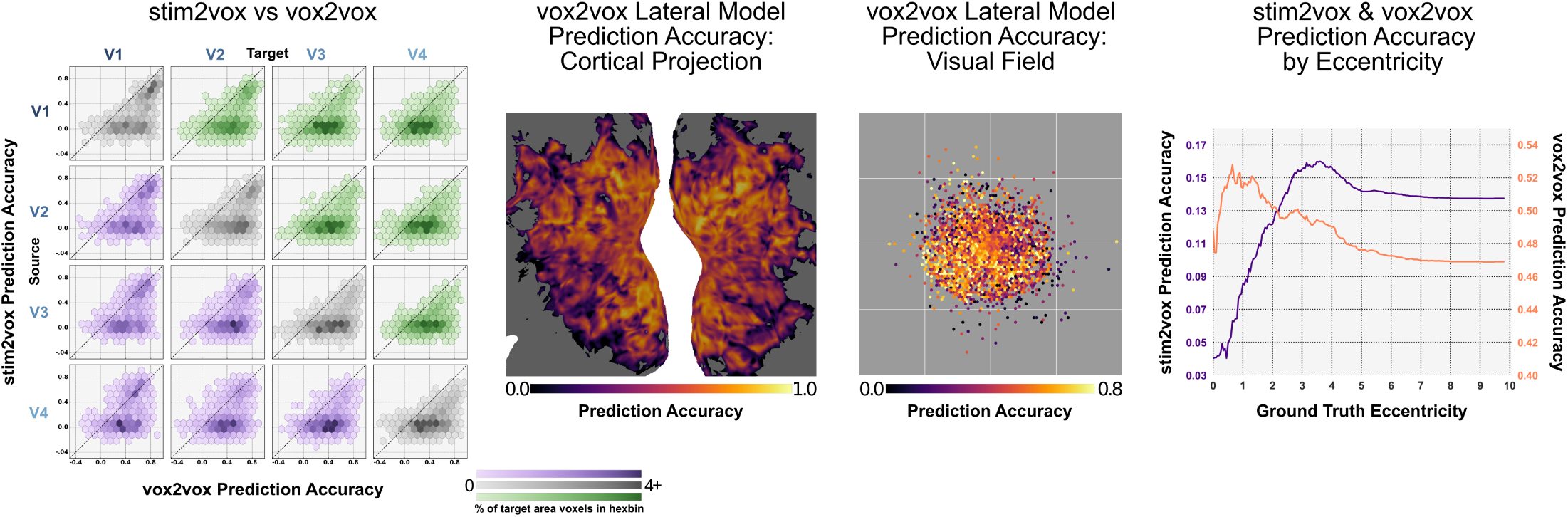
Visualizations of the prediction accuracy of vox2vox models. Should be compared to Figure 1A. **Left** Comparison of prediction accuracy of the stim2vox (y-axis of each panel) and vox2vox models (x-axis) across a matrix of source (rows) and target (columns) visual areas (V1, V2, V3, V4). The vox2vox model generates more accurate predictions than the stim2vox model for most voxels (percentage of voxels in source area indicated by color intensity, light = low, dark = high) for all source/target pairs and all vox2vox model types (green = feed-forward model, gray = lateral models, purple = feedback models). **Middle Left** Prediction accuracy of the lateral vox2vox model (color) projected onto a cortical flatmap. Prediction accuracy is roughly uniform across the cortical surface. **Middle Right** Prediction accuracy of the lateral vox2vox model projected into visual space (format as in Figure 1). Prediction accuracy is roughly uniform across visual space. **Right**: Prediction accuracy against receptive field eccentricity (format as in Figure 1). Prediction accuracy for the vox2vox model (orange, right y-axis) is more uniform across eccentricities and generally higher than predication accuracy for the stim2vox model (purple, left y-axis, data duplicated from Figure 1).

These results show that, for example, the activity in V4 under an optimized linear transformation more accurately predicts activity in V1 than the stimulus under an optimized nonlinear transformation. Thus, the source of the variance that the stim2vox model doesn’t explain is clearly common to many voxels.

### 3.2. Variance unexplained by stim2vox models is subject-specific

We fit linear vox2vox models for source and target voxels in different brains. These cross-subject vox2vox models did not enjoy the dramatic improvement in prediction accuracy over the stim2vox encoding model that we observed when within-subject vox2vox models were applied (Fig. 3). This indicates that the cross-subject vox2vox models are, like stim2vox models, blind to a source of variance that is common to voxels in the same brain.

**Figure 3:**
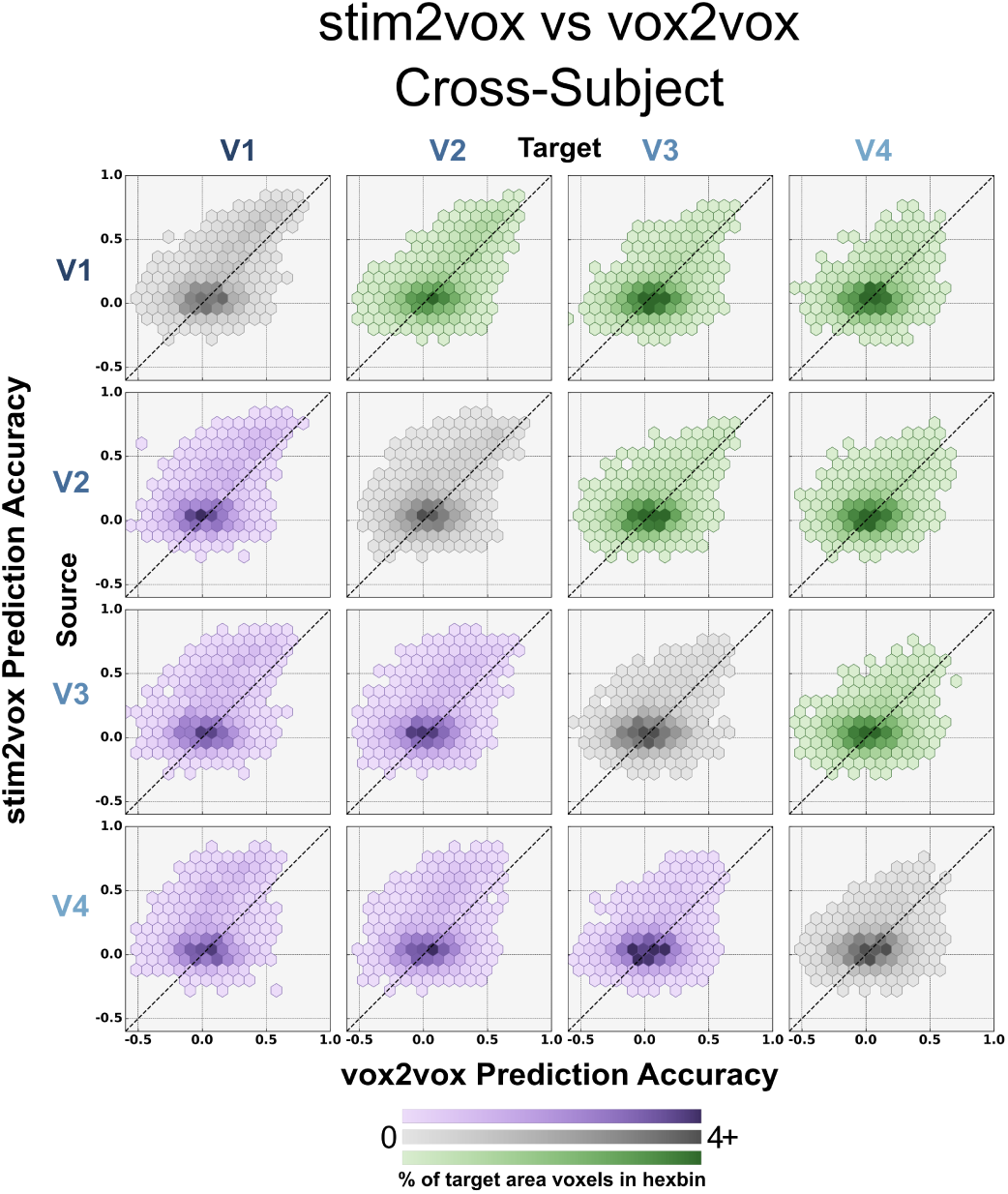
Comparison of stim2vox and cross-subject vox2vox prediction accuracy. Format as in Figure 2. Cross-subject vox2vox models do not enjoy the relative increase in prediction accuracy over stim2vox models as same-subject vox2vox models.

### 3.3. Variance unexplained by stim2vox models is retinotopically mapped

Is the source of variance unexplained in any one voxel common to *all* voxels in the same brain, or only to voxels that have overlapping receptive fields? To answer this question we determined if vox2vox models preferentially connect target voxels to source voxels with receptive fields that overlap the target voxels’. To make this determination we plotted the weights of individual vox2vox models against their receptive field locations. Importantly, we restricted our analysis to target voxels for which the prediction accuracies of the stim2vox and vox2vox models were below and above a common threshold, respectively (Pearson correlation = 0.2; *p* < 0.01, permutation test). In other words, we analyzed only voxels that were “rescued” from the ball of nothingness by their respective vox2vox models. We found that the source voxels with the largest positive vox2vox model weights had receptive field locations that tended to cluster near the receptive field location of the target voxels. On average, the clustering was tight enough that it was possible to accurately estimate the “ground truth” receptive field location (as estimated using a separate retinotopic mapping experiment) of target voxels by simply calculating the center of mass of the receptive field locations of the source voxels with the largest vox2vox model weights (Fig. 4). Although estimates of receptive field location derived from vox2vox models were most accurate when the source and target voxels belonged to the same visual area, estimates were more accurate than receptive field locations derived from the stim2vox model even for hierarchically distant source-target pairings (Fig. 4 top left). Thus, for a given target voxel the source of variance unexplained by the stim2vox models during natural scene stimulation is not shared by all voxels in the same brain, but is shared with (and only with) voxels that have overlapping receptive field locations (i.e., voxels that co-activate during retinotopic mapping stimulation).

**Figure 4:**
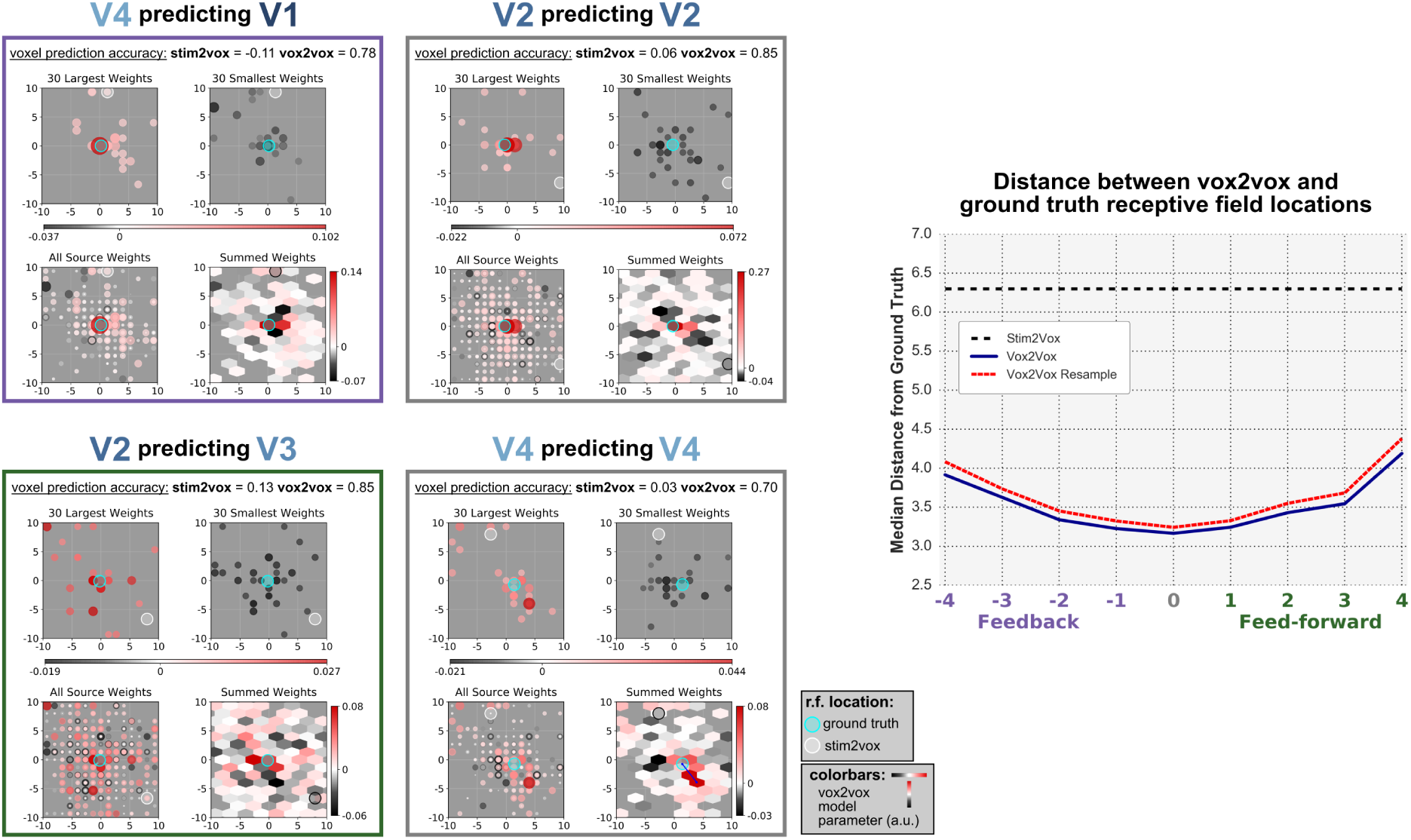
Estimating receptive field location from vox2vox model parameters. **Left**: The parameters of vox2vox models for four example target voxels. For each example voxel the prediction accuracy of the stim2vox model fell below a significance threshold (*p* < 0.01); the prediction accuracy of the vox2vox model was above the threshold. When plotted in visual space (gray panels) according to the receptive field locations of the source voxels, the largest positive vox2vox model parameters (circles in top left of each box; circle radius and intensity scale with magnitude of weight) cluster near the “ground truth” receptive field location of the target voxel (aqua circle; estimated from an independent retinotopic mapping experiment). The largest negative parameters (top right of each box) tend to cluster in the near periphery of the ground-truth receptive field location. The receptive field estimated from the stim2vox model (white circle) is often egregiously misplaced. Visualizations of all parameter values (bottom left of each box) and of local sums of parameter values (bottom right) also reveal distinct peaks of positive parameter values near the ground-truth receptive field location. The receptive field location estimated from the vox2vox model parameters is the location of the peak in the plot of the locally-summed parameter values. The distance between vox2vox and ground truth receptive field location is indicated (blue line) for the V4 to V4 model. **Right**: The median distance (y-axis) between the vox2vox and ground truth receptive field locations (blue curve, red curve is 99^*th*^ percentile of distribution over medians of resampled data) is plotted against hierarchical distance between source and target visual areas. For all source-target pairs this is smaller than the median distance between the stim2vox and ground truth receptive field locations (dashed line).

### 3.4. Prediction accuracy of vox2vox models depends on signed, hierarchical distance between source and target

The relationships between patterns of activity (and the representations those patterns encode) in distinct visual areas in the brain are undoubtedly nonlinear. Intuitively, the relationships between source and target voxels in different brain areas should therefore show some resistance to linear vox2vox modeling. We might expect this resistance to be especially strong for hierarchically distant brain areas that are known to encode stimuli into very different visual features. Thus, we examined median prediction accuracy of the vox2vox models for each pairing of source and target visual area as a function of hierarchical distance and sign.

Consistent with our expectations, we found that median prediction accuracy for any target area was highest for lateral models (i.e., source voxels in same area as area of target voxel) but then declined monotonically as hierarchical distance between a source and target area increased in the feed-forward direction.

Yet several aspects of the relationship between source and target areas were somewhat unexpected. The prediction accuracy of feedback models did *not* decline with hierarchical distance (Fig. 5, S1, S2, A & B) between source and target area, and was higher than the prediction accuracy of the feed-forward model (Fig. 5 S1, S2, D) for most source/target pairs. Finally, while the lateral model was most accurate for each target area, median prediction accuracy for lateral models declined with ascension of the visual hierarchy (Fig. 5 S1, S2, C). These patterns could not be explained by variation in the number of voxels across ROIs, or the percentage of voxels selected as source voxels in each ROI (Fig. S1).

**Figure 5:**
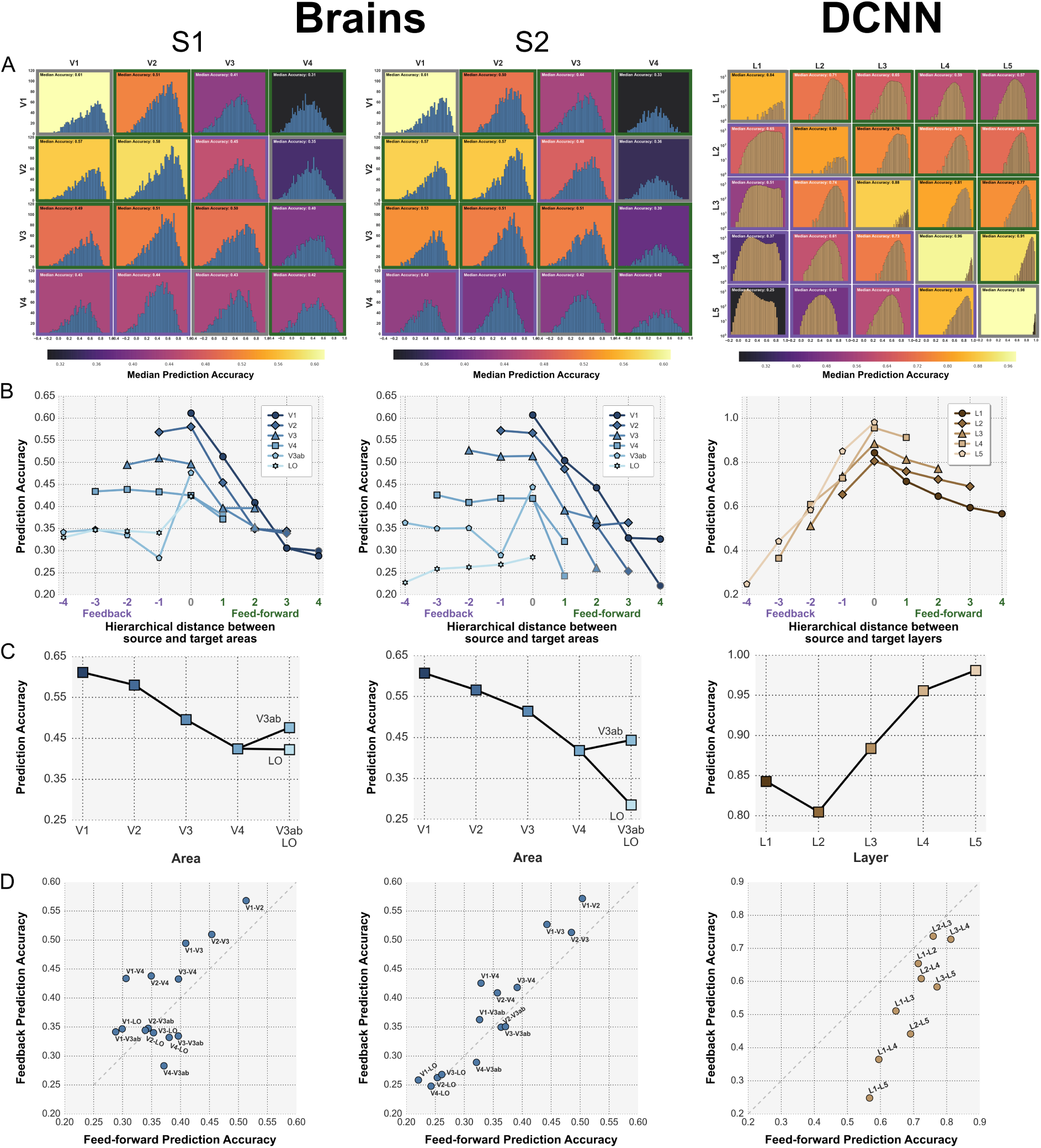
Patterns of prediction accuracy across source-target pairs differs between brains and deep neural networks. **A**: Sub-panels show the distribution (voxel/node count on y-axis) of prediction accuracy (x-axis; background color indicates median of distribution) for vox2vox models (left panels) and pix2pix models (right panel) with the specified source (row) and target (column) pairing. For the brain (left panels) sources and targets are visual cortical regions of interest (ROIs). For the neural network (right panel) sources and targets are layers numbered from L1 (closest to input) to L5 (farthest from input). **B** In the brain median prediction accuracy (y-axis) of feed-forward vox2vox models declines with hierarchical distance (x-axis; 0 indicates lateral model) between source area (indicated by color of each curve) and target area (indicated by distance to source). Median prediction accuracy of feedback vox2vox models is not dependent on hierarchical distance. In the DCNN, median prediction accuracy of feed-forward pix2pix models declines slowly with hierarchical distance; median predication accuracy of feedback models declines more rapidly. **C**: In the brain median prediction accuracy of lateral vox2vox models decreases with hierarchical position of source and target area. In the DCNN median prediction accuracy of lateral pix2pix models increases with hierarchical position of source and target layer. **D**: In the brain median prediction accuracy of feedback vox2vox models (y-axis) is larger than median prediction accuracy of feed-forward vox2vox models (x-axis) for each pair of visual brain areas (blue dots). In the DCNN, median prediction accuracy of feed-forward pix2pix models is smaller than median prediction accuracy of feed-forward pix2pix models for each pair of network layers (brown dots).

Interestingly, a very different relationship between prediction accuracy and hierarchical location was observed when we estimated linear approximations to the connections between layers in the DCNN. We refer to these linear approximations as pixel-to-pixel (pix2pix) models. Like vox2vox models, each pix2pix model predicts activation in a target node of the DCNN as a linear function of activations in a population of source nodes. As in the brain, in the DCNN median prediction accuracy for any target node was highest for lateral models (i.e., source nodes in same layer as the layer of the target node), and median prediction accuracy declined monotonically as hierarchical distance between a source and target layer increased (Fig 5 DCNN, A & B). In contrast to the brain, the prediction accuracy of *feedback* pix2pix model declined *more* rapidly with hierarchical distance between layers in the DCNN than the feed-foward pix2pix models, and was *lower* than the prediction accuracy of the feed-forward model for all source/target pairs. Finally, median prediction accuracy for lateral models *increased* with ascension of the network hierarchy (Fig 5 DCNN, C).

## 4. Discussion

### 4.1. Summary

The central finding of this work (Fig. 2) is that a linear transformation of activity in source voxels (the vox2vox model) predicted activity in nearly every target voxel more accurately than an optimized, nonlinear transformation of the stimulus (the DCNN-based encoding model referred to here as the “stim2vox” model). This finding clearly demonstrates that the stim2vox model is blind to one or more “hidden” sources of variance that induce strong correlations between the activities of voxels across the visual hierarchy. These hidden sources of variance must be endogenous (i.e., not entirely stimulus-dependent) because the vox2vox model did not predict more accurately than the stim2vox model when source and target voxels were located in different brains (Fig. 3). Importantly, we have shown that the correlations induced by these hidden, endogenous sources of variance are highly structured and appear to be dependent upon representations encoded in the brain activity. Induced correlations are strongest between voxels with adjacent receptive fields (Fig. 4 left), even when source and target voxels are hierarchically distant (Fig. 4 right). The extent to which linear vox2vox models can exploit induced correlations to achieve accurate predictions depends upon the hierarchical locations of the source and target voxels (Fig. 5A & B). The prediction accuracy of lateral vox2vox models degrades with ascent of the visual hierarchy (Fig. 5C), and the prediction accuracy of feedback models is larger than the corresponding feed-forward models for most source/target pairs (Fig. 5D). Intriguingly, pix2pix models (i.e., linear approximations of the coupling between units in the DCNN) showed exactly the opposite relationship between prediction accuracy and hierarchical level of the source/target units.

In what follows, we consider in turn a number of possible hidden, endogenous sources of variance that might induce the kind of highly structured correlations observed here. We then consider how we might modify the model of brain computation implied by the DCNN to capture the highly structured variance that the DCNN-based encoding model can’t explain.

### 4.2. Spatially correlated noise

By “noise” we mean measurable variation whose source is not related in any interesting way to what subjects saw or thought during the experiment. There is no question that such noise is present in BOLD measurements, and it is certainly endogenous (i.e., specific to individuals). Furthermore, if such noise were spatially correlated over small distances it might appear to be retinotopically mapped, since the receptive fields of adjacent voxels overlap.

While spatially correlated measurement noise could lead to spurious retinotopic relationships locally, it is unlikely to lead to the kind of long-distance retinotopic registration that was revealed by the weights of the vox2vox models (Fig. 4). Additionally, the asymmetry in prediction accuracy between feed-forward and feedback models (Fig. 5) cannot be accounted for by spatially correlated noise. The feed-forward V1 to V4 model performed much worse than the feedback V4 to V1 model, yet the hierarchical distance is the same. Thus, spatially correlated noise cannot entirely explain our results.

### 4.3. Eye movements

Although subjects were required to maintain fixation on a central dot during the experiment, small involuntary eye movements might effectively translate the stimulus in a random direction on each trial. These random translations could induce endogenous, spatially correlated and even retinotopically mapped variations in activity that would not be captured by a stim2vox model (unless the model somehow incorporated recorded eye movements on each trial; unfortunately, eye movement data is not available for this experiment). This eye-movement-induced variation in activity would most likely be largest in brain areas or regions with small receptive fields and high spatial frequency preference. This would explain why vox2vox models offered a dramatic improvement in prediction accuracy over the stim2vox model for voxels in the foveal representation, and were most effective in low-level visual areas (Fig. 2A and Fig. 5A). Furthermore, eye-movement-induced variation in activity would most likely be smallest in brain areas or regions with large receptive fields and high spatial frequency preference. This would explain why we observed a decrease in vox2vox model prediction accuracy with ascent of the visual hierarchy.

Two aspects of our results challenge the eye-movement interpretation. First, if low-level areas are more influenced by eye-movement than high-level areas, it should be more difficult to predict the activity of target voxels with a feedback model than a feed-forward model, and the difficulty should increase with hierarchical distance below the source voxels. Instead, we observe that the prediction accuracy of feedback models for any source/target pair is almost always greater than for the corresponding feed-forward model, and does not depend upon hierarchical distance. The eye-movement interpretation thus contradicts the feed-forward/feedback asymmetries in prediction accuracy that we observed in our data.

A second challenge to the eye movement explanation is the discrepancy between the results for natural scenes vs. retinotopic mapping experiments. Foveal receptive fields are readily estimated from activity evoked by retinotopic mapping stimuli, but not, as we have shown, by natural scenes. Thus, during retinotopic mapping foveal voxels do not seem to be as dominated by stimulus-independent variance as they are during natural scene stimulation. The fixation task is the same for the retinotopic mapping and natural scenes experiments, so the frequency and magnitude of eye movements during the two experiments is unlikely to differ by much. This discrepancy suggests that the correlations exploited by the vox2vox model may in fact have more to do with the way that natural scenes (as opposed to synthetic stimuli) are processed than with eye movements.

### 4.4. Feature mismatch

The superior prediction accuracy of vox2vox relative to stim2vox models may simply reflect a mismatch between the visual features learned by the DCNN and the native visual features encoded in brain activity. Since vox2vox models accept source voxel activity as input, vox2vox models are obviously better-positioned than stim2vox models to leverage native visual features to predict activity in target voxels.

A challenge to the feature-mismatch interpretation is the inferior prediction accuracy of vox2vox relative to stim2vox models when source and target voxels are in different brains (Fig. 3). This result indicates that the DCNN provides at least as good an estimate of the visual features encoded in the target subject’s brain activity as the brain activity of the source subject. Thus, the visual features “missing” from the DCNN-based stim2vox model would appear to be specific to individual brains. This suggests that the features may be transient and/or dependent on an individual’s idiosyncratic developmental or recent history.

### 4.5. Ongoing activity

A final compelling interpretation of the superior prediction accuracy of vox2vox relative to stim2vox models is that it reflects the well-known predominance of ongoing activity in the visual system [8, 19]. A extensive body of work has shown ongoing, stimulus-independent activity to be meaningfully structured [20, 19], highly correlated across neurons and regions [21], in register with cortical topography [22, 23, 24], and not dismissible as eye movement or noise [24].

Interpreted this way, our results establish that for the vast majority of voxels (i.e., all voxels in the ball of nothingness), and therefore most of visual cortex, ongoing activity is the dominant component of activity measured during vision. Under this interpretation our results further establish that ongoing activity is especially dominant in the foveal representation.

One potential source of ongoing activity that is consistent with its dominance in the foveal representation is top-down feedback emanating from a memory store. Evidence suggests that many forms of top-down signal specifically target foveal representations [25, 26, 27, 28, 29], including top-down signals related to previous experience [30]. Under this interpretation, feed-forward signals evoked by the stimulus may be accurately modeled by the DCNN-based stim2vox model; however, these signals may trigger feedback that is entirely specific to individual experience or memory. Such feedback signals could not be modeled by the DCNN-based stim2vox model, as the DCNN has no explicit representation of feedback or memory.

### 4.6. Toward a better encoding model

As mentioned above, linear vox2vox models cannot possibly provide an adequate characterization of all inter-area relationships in the brain, as the relationships between patterns of activity (and the representations those patterns encode) in distinct visual areas in the brain are undoubtedly nonlinear. Nonetheless they provide more accurate predictions than the DCNN-based encoding model, which is the single best extant stim2vox model. How might the DCNN be substantially altered to make it a more suitable model of the visual system?

A minimum requirement to account for the high prediction accuracy of feedback vox2vox models is that information about low-level representations must somehow be preserved in higher-level activity patterns. For instance, one might imagine a situation where the high-level representation for the picture of a dog contains not only the high-level “dog” object representation but also disentangled representations of various other qualities like fur color, pattern, relative position in space, angle, and so on, that would jointly predict low-level representations. Such disentangling may also account for the decrease in vox2vox lateral model predictability with the ascent of the hierarchy.

The increase in pix2pix lateral model predictability suggests that there remains a high degree of entanglement or within-layer redundancy in the representations of the DCNN used here. The DCNN thus appears to map an entangled high-dimensional image representation into a low-dimensional invariant representation (i.e., an object category). In doing so the network learns to discard “distractor” information in order to produce a very specific subset of invariant representations.

Discriminative models with multiple joint objectives may drive the preservation of information in representations that would otherwise have been discarded as distractors for a single task [31]. Such networks can in principle come closer to achieving the ideal of maximal preservation of information [32] in the process of making it accessible (i.e. disentangled). It would be interesting to see if such models for vision do indeed improve the correspondence between model and brain level-to-level predictability.

## Acknowledgements

This work was supported by grants NIH R01MH111447 and NIH R01EY023384 to TN.

---

**Figure S1:**
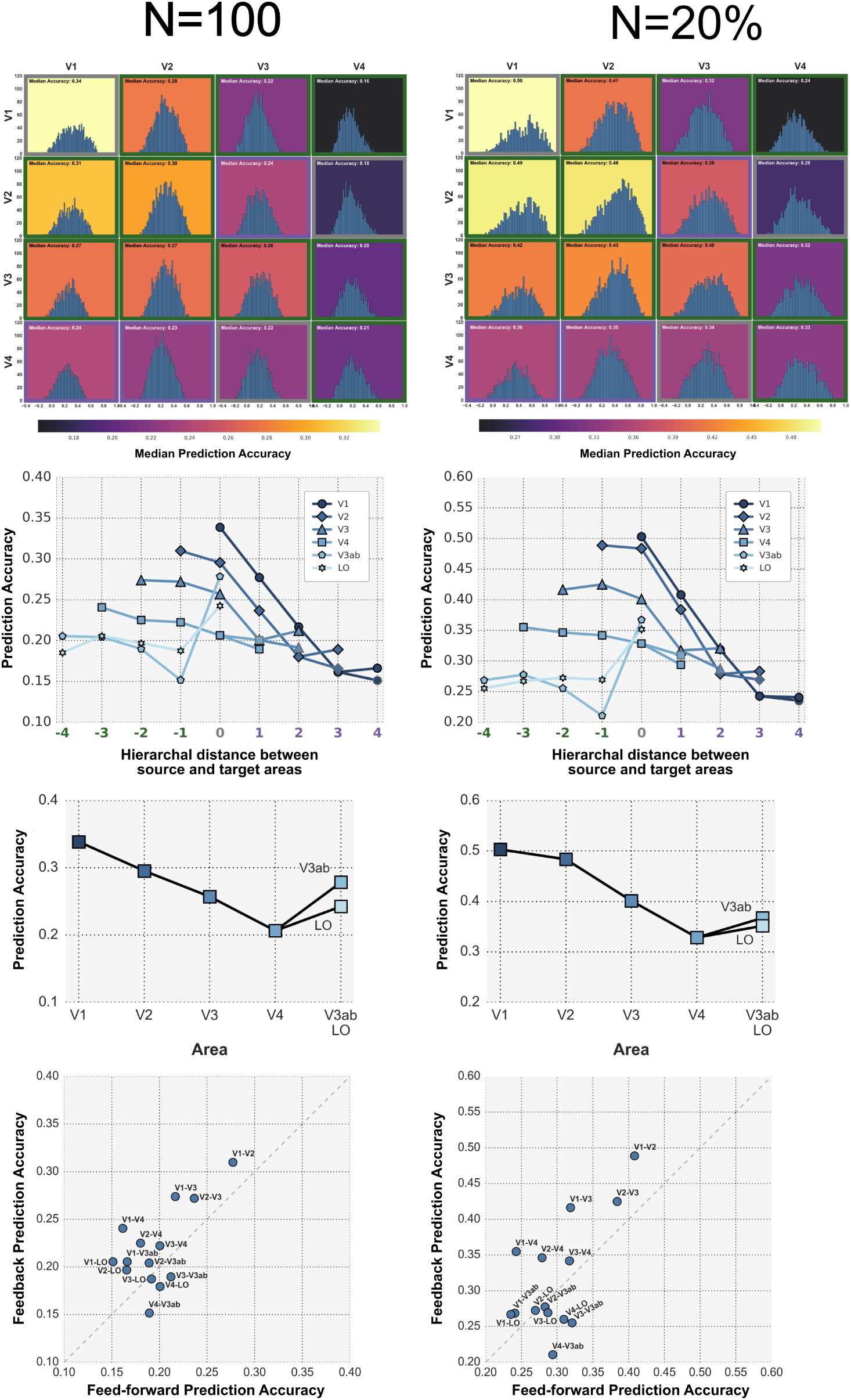
Number of voxels in source model doesn’t change prediction accuracy pattern. In order to determine if prediction accuracy was influenced by the differing number of voxels in each area, we employed two sampling strategies. First we randomly sampled 100 source voxels (A), 10 times for each source-target pairing. We averaged across the 10 iterations and plotted in the same way as Figure 5. Next we randomly sampled 20% (B) of source voxels and proceeded in the same manner. In both sampling strategies, the same pattern of declining median prediction accuracy in feed-forward and lateral directions and stable accuracy in feedback directions was observed. All data from S1.

